# Do transformers and CNNs learn different concepts of brain age?

**DOI:** 10.1101/2024.08.09.607321

**Authors:** Nys Tjade Siegel, Dagmar Kainmueller, Fatma Deniz, Kerstin Ritter, Marc-Andre Schulz

## Abstract

“Predicted brain age” refers to a biomarker of structural brain health derived from machine learning analysis of T1-weighted brain magnetic resonance (MR) images. A range of machine learning methods have been used to predict brain age, with convolutional neural networks (CNNs) currently yielding state-of-the-art accuracies. Recent advances in deep learning have introduced transformers, which are conceptually distinct from CNNs, and appear to set new benchmarks in various domains of computer vision. However, transformers have not yet been applied to brain age prediction. Thus, we address two research questions: First, are transformers superior to CNNs in predicting brain age? Second, do conceptually different deep learning model architectures learn similar or different “concepts of brain age”? We adapted a Simple Vision Transformer (sViT) and a Shifted Window Transformer (SwinT) to predict brain age, and compared both models with a ResNet50 on 46,381 T1-weighted structural MR images from the UK Biobank. We found that SwinT and ResNet performed on par, while additional training samples will most likely give SwinT the edge in prediction accuracy. We identified that different model architectures may characterize different (sub-)sets of brain aging effects, representing diverging concepts of brain age. Thus, we systematically tested whether sViT, SwinT and ResNet focus on different concepts of brain age by examining variations in their predictions and clinical utility for indicating deviations in neurological and psychiatric disorders. Reassuringly, we did not find substantial differences in the structure of brain age predictions between model architectures. Based on our results, the choice of deep learning model architecture does not appear to have a confounding effect on brain age studies.

## 1. Introduction

The brain undergoes structural changes while aging (MacDonald and Pike, 2021), accompanied by reduced cognitive function and increased risk of neurodegenerative disorders (Peters, 2006; Farooqui and Farooqui, 2009). The rate at which aging alters the brain appears to be influenced by the presence of disease (Anderton, 1997), lifestyle (Peters, 2006), and environmental factors (Esiri, 2007).

*Brain age prediction* estimates biological age using machine learning (ML) techniques applied to neuroimaging data. Such brain age prediction models are generally trained on healthy cohorts, ensuring that the model learns the amount of aging considered normal for healthy subjects (Feng et al., 2020; Dinsdale et al., 2021; Kolbeins-son et al., 2020). Differences between brain-predicted age and chronological age (brain age gap, BAG) (Ballester et al., 2023; Chen et al., 2022; Man et al., 2021) have been elevated for patients with various psychiatric and neurological disorders, including Alzheimer’s disease (AD), Parkinson’s disease (PD), multiple sclerosis (MS), mild cognitive impairment (MCI), major depression (MD), schizophrenia and bipolar spectrum disorder (BSD) (Beheshti et al., 2020; Cole et al., 2020; Eickhoff et al., 2021; Bashyam et al., 2020; Nenadić et al., 2017; Kaufmann et al., 2019). Elevated BAGs have also been linked to markers of poor health such as obesity, high blood pressure, and diabetes (Wrigglesworth et al., 2021). This elevation in BAGs is thought to arise from an overlap between the effects of aging, the secondary neurobiological effects of diseases and poor general health (Cole and Franke, 2017). The accumulating evidence linking BAGs with various health-related factors and neurological and mental diseases has boosted the popularity of BAGs as individualized biomarkers of structural brain health (Cole and Franke, 2017).

It is commonly believed that accurate brain age models are essential to provide useful biomarkers (Hahn et al., 2021; Peng et al., 2021; Cole, 2020; Tanveer et al., 2023; Niu et al., 2020), and deep learning with convolutional neural networks (CNNs) has yielded the most accurate age predictions to date (Peng et al., 2021; Gong et al., 2021; Leonardsen et al., 2022). Deep learning models such as CNNs are capable of operating on minimally processed neuroimaging data, primarily voxel-wise structural magnetic resonance imaging (sMRI) brain images (Feng et al., 2020; Dinsdale et al., 2021; Lee et al., 2022; Peng et al., 2021; Leonardsen et al., 2022). Using voxel-wise input images, CNNs can learn to model complex visual features of brain aging from the ground up.

A recent innovation in deep learning architectures has been the development of transformer models Vaswani et al. (2017), such as vision transformers (Dosovitskiy et al., 2021). Whereas single layers of CNNs combine information that is locally related in the input, vision transformers are more flexible. They can create visual features by including information from different parts of the input, even if these are not spatially close together. Although the greater flexibility of vision transformers comes at the cost of requiring substantially larger amounts of training samples (Dosovitskiy et al., 2021), vision transformers seem to surpass the CNN set benchmarks in various domains of computer vision, including image classification (Dosovitskiy et al., 2021), semantic segmentation (Xie et al., 2021), and object detection (Liu et al., 2022). Considering the success of vision transformers naturally prompts the question: Can transformers be utilized to make brain age predictions more accurate? And - since it is conceivable that characterizing only a small subset of aging effects in the brain is sufficient for accurately predicting age - do conceptually distinct deep architectures learn different “concepts of brain age” (see Section 3.1)? As the mechanism by which CNNs and transformers process input data fundamentally differs, CNNs for brain age predictions could learn to characterize one subset of brain aging effects, while transformers could learn to characterize another.

If different deep learning model architectures attend to different concepts of brain age, this would have profound implications for the brain age research paradigm. First, different model architectures could confound the results of prior studies, because different concepts of brain age could identify different disease-related patterns. Comparing how informative BAGs are to diseases and health related factors would become challenging if different model architectures are employed, even in similar cohorts. Second, selecting a model architecture for brain age prediction would become increasingly complicated. For instance, one brain age concept could encompass a broad range of disease-patterns, while others entail only few. Hence, the selection of a model architecture would require measures of clinical utility rather than solely relying on model accuracy, which is the current common practice (Han et al., 2022; Baecker et al., 2021; Niu et al., 2020; Kuo et al., 2021; Amoroso et al., 2019). Third, if different brain age concepts inform on specific diseases, the role of BAGs as general brain health biomarkers, previously highlighted Cole and Franke (2017), would require reevaluation. Practically, identifying which model architecture corresponds to which brain age concept would be essential, as BAGs might indicate specific diseases rather than general brain health.

To investigate whether different deep learning model architectures learn different concepts of brain age and whether a transformer can predict brain age with superior accuracy, we adapted the popular simple vision transformer (sViT) (Beyer et al., 2022) and shifted window transformer (SwinT) (Liu et al., 2021) to predict age from 3D T1-weighted sMRI brain scans. For comparison, we trained a ResNet He et al. (2016), which is one of the most widely used CNNs in brain age prediction (Fisch et al., 2021; Jónsson et al., 2019; Kolbeinsson et al., 2020; Ballester et al., 2021; Shah et al., 2022; Hu et al., 2023). We systematically investigate whether ResNet, sViT, and SwinT attend to different concepts of brain age by examining differences in their predictions and clinical utility (ability to inform neurological and psychiatric diseases, health-related factors; see Section 3.2) as proxies for differences in how “brain age” is characterized by either model architecture (see Section 3.3). Divergent predictions and clinical utility across model architectures would be indicative of variations in the model architectures’ concepts of brain age. To measure clinical utility we concentrate on diseases commonly examined in brain age studies, namely PD (Eickhoff et al., 2021), MS (Cole et al., 2020), epilepsy (Sone et al., 2021), alcohol use disoder (AUD) (Bøstrand et al., 2022), bipolar affective disorder (BAD) (Hajek et al., 2019), and psychotic disoders (Ballester et al., 2022)), as well as factors associated with brain health, specifically fluid intelligence, reaction time, trailmaking interval (Smith et al., 2019), tobacco consumption (Franke et al., 2013), mobile phone usage (Thomée, 2018), TV consumption (Dougherty et al., 2022), systolic blood pressure (Smith et al., 2019), grip strength (Carson, 2018), and body mass index (BMI) (Ward et al., 2005). An overview of our workflow and results is displayed in Figure 1.

**Figure 1:**
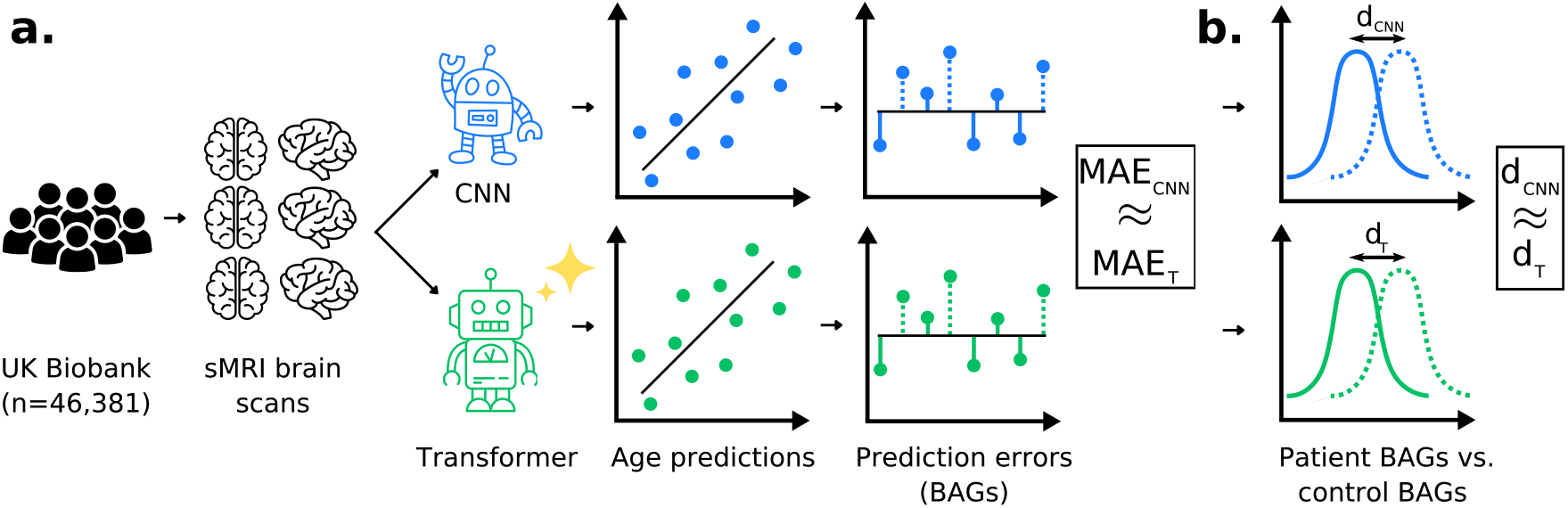
Overview of workflow and results: **a**. We used 46.381 structural magnetic resonance imaging (sMRI) brain scans from the UK Biobank to train and evaluate a convolutional neural network (CNN; 3D ResNet50) and two transformers (3D simple vision transformer; sViT; 3D shifted window transformer; SwinT) for brain age prediction. Mean absolute errors (MAEs) for held-out healthy subjects were nearly identical for ResNet (2.66 years) and SwinT (2.67 years). Extrapolating MAEs beyond the number of available brain images revealed that SwinT will most likely become more accurate than ResNet with increased training samples. **b**. Effect sizes between prediction errors (brain age gaps; BAGs) of patients and matched controls were similar for CNN and transformers across neurological- and psychiatric diseases, yielding no indication that different model architectures rely on different brain aging effects for their predictions.

## 2. Related work

Previous works have been concerned with technical aspects of brain age prediction, such as bias correction (Beheshti et al., 2019; de Lange and Cole, 2020; Zhang et al., 2023; Liang et al., 2019), performance metrics (de Lange et al., 2022), and prediction accuracy of different ML models (Valizadeh et al., 2017; Baecker et al., 2021; Lam et al., 2020). Other brain age model comparisons additionally included reliability measures (Bacas et al., 2023; Dörfel et al., 2023), aggregate measures of clinical utility (Lee et al., 2021; More et al., 2023; Xiong et al., 2023; Lee, 2023; Beheshti et al., 2021), and general feature importance (Ball et al., 2021; Han et al., 2022). Also, there are brain age studies that have investigated general feature importance for CNNs (Lee et al., 2022; Hepp et al., 2021; Levakov et al., 2020; Hofmann et al., 2022).

To the best of our knowledge, no previous work explicitly considers the possibility that the fundamental “concept of brain age” could mismatch between ML models, nor do any of the studies include deep learning model architectures that are conceptually different.

In addition, we believe to be the first to present cutting-edge transformers for brain age prediction from sMRI data. So far, transformers have solely been combined with CNN-based feature encoders, to fuse information from different image scales (He et al., 2021a) and modalities (Zhao et al., 2024; Cai et al., 2022; He et al., 2021b), to punctually add global information pathways (Hu et al., 2022), or to refine CNN-extracted features from 2D image slices (Jun et al., 2021).

## 3. Theory

To clarify our study, we introduce three key concepts. First, we redefine the “concept of brain age”, suggesting the existence of multiple brain ages, necessitating more precise terminology. Second, we discuss the “clinical utility” of these concepts, aiming to quantify their usefulness in clinical settings. Third, we explore latent representations in brain age models that encode these varied concepts and discuss methods to probe these representations.

### 3.1. Different concepts of brain age

In the past, brain age has generally been regarded as a uniform concept, yet different models may accurately predict age while relying on different brain aging effects. Such aging effects may include loss of total brain volume, enlargement of ventricles, cortical thinning (especially in frontal areas) and shrinkage of subcortical gray matter structures, with specific structures (e.g. hippocampus) shrinking at increased rate (MacDonald and Pike, 2021). To distinguish different combinations of brain aging effects, we introduce the term “concept of brain age”, referring to the features models use for age prediction (e.g. ventricle size and frontal lobe thickness), and how these features are combined (e.g. ventricle size prioritized over frontal lobe thickness).

Brain age concepts may vary in the particular features they consider as well as in the number of features they consider, reflecting uncertainty about whether a broad or narrow range of aging indicators is necessary for accurate predictions. This uncertainty stems from the fact that different brain features carry redundant information on age. For example, Bethlehem et al. (2022) have computed detailed normative trajectories for various brain morphological features across the human lifespan. Each of those trajectories, or combinations, could, in theory, be learned by (non-linear) models to predict age. Which particular brain features are picked up by a given model may depend on factors such as model architecture, initialization, amount of training data, and model capacity - in addition to the relationship between brain features and biological aging.

One might argue that different concepts of brain age merely refer to different ways of measuring “brain age”. However, considering that disease and brain health-related factors exhibit regional predilections (Geng et al., 2006; Raz and Rodrigue, 2006; Dekker et al., 2021; Gómez-Apo et al., 2021; Gallinat et al., 2006), it becomes clear that brain age concepts really define brain age’s nature. For example, hypertension appears to accelerate hippocampus shrinkage (Raz et al., 2005), and hence, models with brain age concepts based on hippocampal volume will likely show increased brain age for hypertension. In contrast, models based on brain features that are little affected by hypertension may show no such effect. Ideally, brain age concepts would cover holistic sets of brain aging features, but due to brain features carrying redundant information on aging (Bethlehem et al., 2022), it is questionable whether current brain age models learn such comprehensive brain age concepts.

### 3.2. Clinical utility

Evaluating the practical usefulness of brain age concepts requires a benchmark measure. To that end, we define “clinical utility” as the ability of a brain age model to inform on a broad range of diseases and health-related phenotypes. Specifically, we evaluate “clinical utility” in two ways: first, by examining the sensitivity of BAGs to differences between healthy individuals and those with neurological and mental disorders (Cole et al., 2020; Bashyam et al., 2020; Kaufmann et al., 2019); second, by evaluating how predictive BAGs are of health-related phenotypes (Cole, 2020; Steffener et al., 2016; Lee, 2023).

### 3.3. Probing differences in model architectures’ latent concepts of brain age

We aim to determine if different deep learning model architectures yield distinct brain age concepts. Due to the complex, non-linear nature of these models, it is challenging to identify the features they use for predictions (Kindermans et al., 2019; Adebayo et al., 2018; Sundararajan et al., 2017; Hooker et al., 2019; Dombrowski et al., 2019). Here, we propose using “clinical utility”, i.e., the models’ predictions and prediction errors as proxies for understanding potential differences in latent brain age concepts.

Analyzing the sensitivity of prediction errors (BAGs) for brain diseases and health-related phenotypes provides insights into differences in brain age concepts, as these conditions show a tendency to affect specific brain regions. For example, hypertension has been linked to accelerated hippocampus shrinkage (Raz et al., 2005), PD patients have shown significant pallidum volume loss Geng et al. (2006), tobacco use has appeared to reduce gray matter volume and density in frontal, occipital, and temporal lobes Gallinat et al. (2006), MS has been associated with cerebellar and thalamic atrophy alongside white matter lesions (Dekker et al., 2021), and obesity has been related to gray matter reduction in frontal and temporal regions, basal nuclei, and cerebellum (Gómez-Apo et al., 2021). Thus, one brain age concept might address disease- or behavior related alterations, while another may not.

## 4. Material and methods

### 4.1. Participants

Our study is based on the UK Biobank (UKBB), which is an ongoing prospective biomedical data collection initiative (Sudlow et al., 2015). Specifically, we used data from 46,381 individuals (53% female, age range 44-83, age mean 64.26, age standard deviation 7.75), for whom T1-weighted sMRI brain scans were available at the time of writing. We divided subjects into a normative cohort with no diagnoses in ICD-10 category F (mental and behavioral disorders) and G (diseases of the nervous system), and a patient cohort including all diagnosis in category F and G. To determine how sensitive BAGs are to neurological and psychiatric disorders, we focus on disorders that are frequently studied in the context of brain age research, specifically patients with Parkinson’s disease (PD) (Eickhoff et al., 2021), multiple sclerosis (MS) (Cole et al., 2020), epilepsy (Sone et al., 2021), alcohol use disorder (AUD) (Bøstrand et al., 2022), bipolar affective disorder (BAD) (Hajek et al., 2019) and psychotic disorders^1^ (Ballester et al., 2022) in conjunction with controls from the normative cohort. We selected controls by matching normative subjects to the disease cohorts for each diagnosis using propensity score matching, while controlling for sex, age, education level, household income, the Townsend deprivation index, and genetic principal components, as described in (Schulz et al., 2024b). The remainder of the normative cohort was used for model training. Patients who were not used to measure BAGs’ sensitivity to diseases (patients were also not used for model training) were used to validate the hyperparameters of the model architectures, which led to the following set sizes: *n*_*train*_ = 27, 538, *n*_*val*_ = 16, 499, *n*_*control*/*test*_ = 1, 172.

### 4.2. sMRI data

We used minimally preprocessed 1mm T1-weighted sMRI brain scans provided by the UKBB. The images were skull-stripped with the UKBB-provided brain mask, linearly registered on MNI152 with the UKBB-provided transformation matrices, and center-cropped, resulting in a final resolution of 160×192×160. Preprocessing is in line with literature defaults (Peng et al., 2021; Leonardsen et al., 2022; Fisch et al., 2021; Kolbeinsson et al., 2020).

### 4.3. Target phenotypes

In addition to the sMRI data, we used phenotypic data from the UKBB. Specifically, the UKBB provides information on ICD-10 diagnosis in terms of first occurrence dates, and we assigned disease labels if the first occurrence date was before the date on which the sMRI data were collected. The mappings from diseases to UKBB fields are shown in the Supplement Table S1. To analyze the informativeness of the BAGs for various brain health-associated factors, we used UKBB variables for cognitive performance (fluid intelligence, reaction time, trailmaking interval; Smith et al. 2019), lifestyle choices (tobacco consumption; Franke et al. 2013, mobile phone usage; Thomée 2018, TV consumption; Dougherty et al. 2022) and biomedical condition (systolic blood pressure; Smith et al. 2019, grip strength; Carson 2018, body mass index; Ward et al. 2005). The mapping of each variable to the UKBB field number is provided in the Supplementary Table S2.

### 4.4. Deep learning model architectures

#### 4.4.1. 3D ResNet50

As CNN architecture, we used a ResNet50 (He et al., 2016), adapted to 3D input (Hara et al., 2018). ResNet is a well known standard architecture in computer vision and is widely used in brain age prediction (Fisch et al., 2021; Jónsson et al., 2019; Kolbeinsson et al., 2020; Ballester et al., 2021; Shah et al., 2022; Hu et al., 2023). Conceptually, a simpler form of the ResNet is the VGG (Simonyan and Zisserman, 2025) (or in its shallow form, the SFCN; Peng et al. 2021), which some brain age studies employ, too (Tanveer et al., 2023). In brief, the main component of ResNet (and VGG) is the convolutional layer, which incorporates convolutional filters that slide across the input image and combine local image information to create visual features such as edges or shapes. In our experiments, we used a conventional PyTorch implementation^2^ of the 3D ResNet50 (Hara et al., 2018), with a total number of 46.2 million trainable parameters.

#### 4.4.2. 3D simple vision transformer

In contrast to CNNs, which combine local image information using convolutional filters, vision transformers (Dosovitskiy et al., 2021) process images through a different mechanism. Essentially, vision transformers divide the input image into a sequence of image patches, and then combine information across these patches to characterize visual features. Since all image patches are connected to each other through a so-called attention mechanism (Vaswani et al., 2017), vision transformers can generate visual features composed of spatially unrelated information in the input image. In contrast, CNNs are limited to combining information from local image neighborhoods to form visual features.

In this study, we adapted an sViT (Beyer et al., 2022) to predict age from 3D sMRI scans. A brief description of the specific modifications is given in Appendix A. The 3D sViT implementation we used can be found in the GitHub repository vit-pytorch^3^. Hyperparameters were kept at the vit-pytorch defaults. The complete set of hyperparameters is shown in the Appendix Table A1, resulting in a total of 42.0 million trainable parameters.

#### 4.4.3. 3D shifted window transformer

The SwinT (Liu et al., 2021) is another modification of the original vision transformer (Dosovitskiy et al., 2021), which reintroduces core properties of CNNs to improve performance on visual tasks. In brief, the SwinT divides input images into image patches like the vision transformer. However, it focuses on forming visual features by combining information from locally related image patches, while distant image patches are only connected via indirect pathways. This modification means that the SwinT loses some of the vision transformer’s flexibility in creating visual features, but large images in particular can be processed more efficiently. In addition, the SwinT fuses image patches at different levels of depth, which makes the SwinT learn hierarchical image representations, which have appeared to be crucial for biological vision (Hubel and Wiesel, 1962), and are an essential property of CNNs (LeCun et al., 2010).

Similar to the sViT, we adapted the SwinT to operate on 3D input (Appendix A). Our implementation and hyperparameter choices regarding the number of attention heads, patch size, embedding dimension, and attention window size were based on the SwinUNETR model (Hatamizadeh et al., 2021), which has been used for 3D brain tumor segmentation. The hyperparameter choices for the depths of the model and the expansion ration of the multilayer perceptron (MLP) α were inspired by the “Swin-T” model variant proposed by Liu et al. (2021). The complete list of hyperparameters used for the SwinT model is shown in the Appendix Table A2, which resulted in a total of 10.1 million trainable parameters.

### 4.5. Model training

All model architectures were trained using the PyTorch Lightning 1.8 interface for PyTorch 1.12 and a single Nvidia A100 GPU with 80GB memory for ResNet and sViT, and two A100s of the same type for the SwinT. Each model was optimized using Adam (Kingma and Ba, 2014) on the mean squared error loss, with a one-cycle learning rate policy (Smith and Topin, 2019; Fisch et al., 2021; Schulz et al., 2022). The maximum learning rate for SwinT and sViT was set to 10^−4^ and to 10^−2^ for the ResNet. The training duration was 150,000 gradient update steps for each model architecture. The effective batch size was 8 for ResNet and SwinT, and 16 for sViT. Each model architecture was re-trained 6 times with different random initialization and batch order.

### 4.6. Measuring clinical utility

We measured the clinical utility of BAGs by how sensitive they were to neurological and mental diseases (AD, PD, MS, depression, schizophrenia, BSD), as well as by how predictive they were for health-related phenotypes (fluid intelligence, reaction time, trail making interval, tobacco consumption, mobile phone usage, TV consumption, systolic blood pressure, grip strength, and body mass index). In detail, our analysis workflow proceeded as follows (see Figure 1 for an overview): We first trained multiple instances of sViT, SwinT and ResNet using the normative cohort. Next, we computed BAGs of held-out patients and controls by subtracting chronological age from predicted age, for each of the models’ instances. Then, we measured how sensitive the BAGs were to diseases by calculating effect sizes (Cohen’s *d*) between the BAGs of the patients and matched controls. In addition, we estimated the uncertainties of effect sizes across patient-control pairs via bootstrapping. Next, we analyzed how predictive BAGs were for various health-related phenotypes, by fitting linear models from BAGs and covariates (age, sex, genetic principal components 1-3, years of education, income level) to the phenotypes. For each resulting linear model, we report the t-statistic for the BAG’s β-coefficient, as a measure of how predictive BAG is for the phenotype in question. Again, uncertainties were estimated via bootstrapping.

### 4.7. Measuring consistency of brain age concepts across train runs

Varying brain age concepts could arise because of differences in model architecture, but also due to different random weight initialization and batch order during training. To investigate, we trained 6 instances of each model architecture with varying initializations and batch orders. We analyzed how correlated predictions of either model architectures’ instances were for held-out patients and controls using Pearson’s correlation coefficient, to gain an understanding of potential differences in brain age concepts across train runs, following the same logic as described in Section 3.3.

## 5. Results

### 5.1. SwinT is competitive and will likely outperform ResNet with increasing sample sizes

To investigate whether transformers may outperform CNNs in accurately predicting brain age, we compared mean absolute errors (MAEs) for held-out healthy subjects between SwinT, sViT, and ResNet. SwinT (MAE of 2.67 ± 0.02, mean and SD over different train runs) and ResNet (MAE of 2.66 ± 0.05) performed on par (Table 1). sViT performed noticeably worse, with an averaged MAE of 3.02 ± 0.08 years.

**Table 1:**
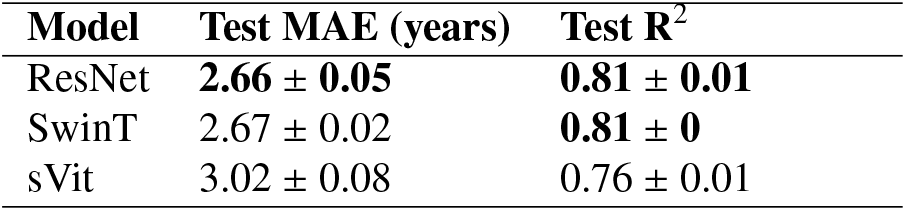
SwinT is competitive to ResNet in brain age prediction. Mean absolute errors (MAEs) and coefficient of determination (R^2^) are displayed for the held-out set of healthy subjects (*n*=1172). The uncertainty estimates indicate the standard deviation (SD) across different randomly initialized model instances. 3D ResNet50 (ResNet) and 3D shifted window transformer (SwinT) predict age with nearly identical accuracy, both outperforming the 3D simple vision transformer (sViT).

In addition, we analyzed how each model architecture’s accuracy scales with the number of training samples. In short, we trained instances of each model architecture with stepwise reduced training samples, and assumed a power-law relation between accuracy and the amount of data (Schulz et al., 2024a). This enabled us to extrapolate each model architecture’s accuracy beyond the amount of training samples available to us. We found that the SwinT can be expected to outperform the ResNet starting from approximately *n* = 25, 000 samples (Figure A1), with the ResNet marginally benefitting from more training samples. The sViT’s performance can be expected to benefit from increasing training samples, though it may not be able to achieve accuracies comparable to SwinT and ResNet in its current form.

### 5.2. No evidence that sViT, SwinT and ResNet attend to different concepts of brain age

To investigate whether SwinT, sViT and ResNet may attend to different concepts of brain age, we analyzed differences in predictions and prediction errors as proxies of differences in the underlying aging characterizations (see Section 3.3). In a first analysis, we computed the Pearson correlation for held-out-set predictions between model architectures. Predictions from all three model architectures were highly correlated (average correlation with SD between predictions of differently initialized SwinT and ResNet instances: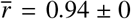, SwinT-sViT: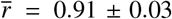, ResNet-sViT: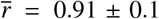, suggesting that each model architecture follows a similar concept of brain age.

In a second analysis, we compared the clinical utility (Section 3.2) of each model architectures’ BAGs. Deviations in clinical utility between model architectures would hint to differences in the concepts of brain age (see Section 3.3). We found that the sensitivity of BAGs for the investigated disorders were comparable across model architectures. Patients’ BAGs were elevated for each model architecture and disease (Figure 2). We observed (in Cohen’s terminology; Cohen, 2013) small effects for epilepsy, small to medium effects for PD, AUD, BAD and psychotic disorders, and medium to large effects for MS. Effect sizes between model architectures were within one σ from each other for any disease, with no indication of differences. The association of BAG and cognitive, lifestyle, and biomedical phenotypes was also comparable across model architectures. Again, the measured effects were within approximately one σ from each other, again with no indication of a difference (Figure 3).

**Figure 2:**
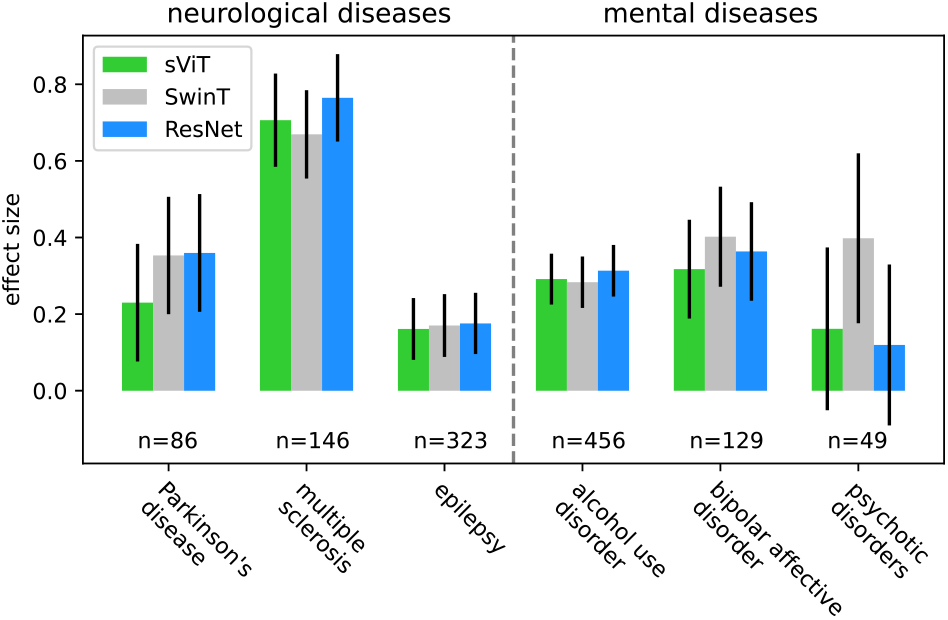
Different brain age model architectures encode similar disease patterns. The figure shows effect sizes (Cohen’s *d*) measured between BAGs of patients and matched controls. Effect sizes between model architectures were within one σ from each other for any disease, with no indication of differences. Error bars indicate the standard error of the mean estimate derived by bootstrapping patient-control pairs.

**Figure 3:**
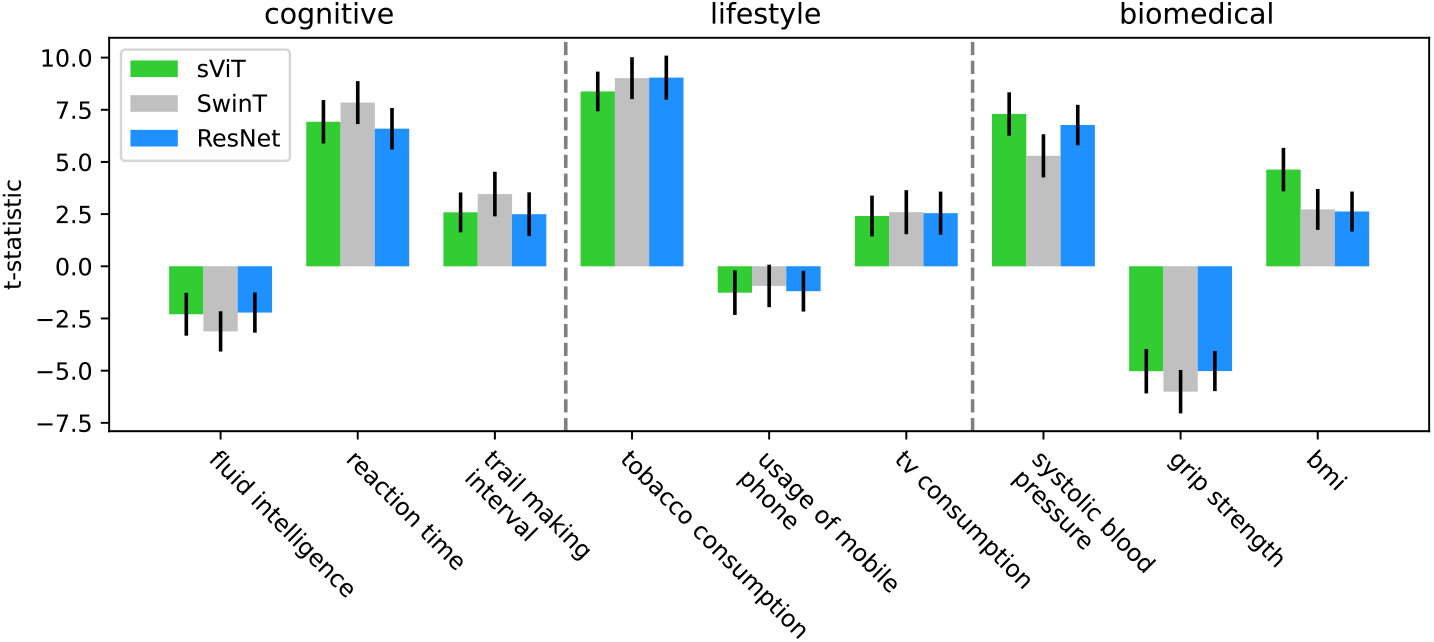
Association of BAG and cognitive, lifestyle and biomedical phenotypes seems not to depend on the model architecture. We fitted linear models from BAG and confounds to phenotype and report the t-statistic for whether the BAG is a significant predictor. Error bars indicate the t-statistic’s standard error of the mean estimate, derived by bootstrapping. BAGs of different model architectures were similarly predictive for the analyzed phenotypes.

The size and directionality of effects was compatible with literature expectations: Weak results on cognitive tests, unhealthy habits, and markers of poor physical condition were associated with elevated BAG, while good results on cognitive tests and markers of good physical condition were associated with a decreased BAG (Smith et al., 2019). In sum, our results suggest that sViT, SwinT and ResNet most likely do not attend to meaningfully different concepts of brain age; we found that predictions of each model architecture were highly correlated, that BAGs were similarly sensitive to neurological and psychiatric diseases, as well as comparably predictive for cognitive, lifestyle, and biomedical phenotypes.

### 5.3. Concepts of brain age appear consistent across train runs

To assess the consistency of either model architecture’s brain age concept across random initializations and batch orders, we computed correlations between held-out-subject predictions within each model architecture and found no indication of varying brain age concepts. Over six different train runs, SwinT averaged a Pearson correlation of 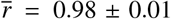 (SD) 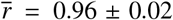 for ResNet; 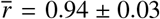 for sViT), suggesting that brain age concepts are mostly unaffected by random initializations and batch order. In comparison to sViT and ResNet, the SwinT appears to converge to more uniform brain age concepts.

## 6. Discussion

In the present study, we make three central contributions. First, we adapt and evaluate the recently popularized transformer architecture for brain age prediction. Using one of the largest brain imaging datasets currently available, we found that the novel SwinT and the widely used ResNet predict age with nearly identical accuracy.

Our results indicate that both evaluated transformer architectures will benefit from growing sMRI datasets, while the accuracy of ResNet appeared to be saturated. Second, we identify that “brain age” might not refer to a uniform concept and outline why “concepts of brain age” may differ between brain age models. Third, we investigate whether conceptually different deep learning model architectures attend to different concepts of brain age. We probed the model architectures for structural differences in their brain age predictions under a range of neurological and psychiatric disorders and with regard to biomedical, cognitive, and behavioral phenotypes, but found no indication that SwinT, ResNet, and sViT may attend to different concepts of brain age.

### 6.1. Transformers for accurate brain age prediction

A common belief in the brain age community is that models need to accurately predict age, in order to provide useful biomarkers (Hahn et al., 2021; Peng et al., 2021; Cole, 2020; Tanveer et al., 2023; Niu et al., 2020). Thus, research on more accurate brain age models has been a dominant topic in the brain age literature, and best accuracies were achieved by CNNs (Peng et al., 2021; Gong et al., 2021; Leonardsen et al., 2022). A more recently popularized deep learning model architecture is the vision transformer, which apparently surpasses benchmarks set by CNNs in various tasks of computer vision (Dosovitskiy et al., 2021; Xie et al., 2021; Tu et al., 2022). Consequently, the question arises whether transformers can be leveraged to obtain more accurate brain age predictions. We applied two of the most popular vision transformers to brain age prediction, and found that the SwinT matches, and for larger sample sizes likely outperforms (Figure A1) the widely used ResNet CNN in brain age prediction accuracy (ResNet MAE 2.66 years, SwinT MAE 2.67). Other works that trained CNNs on UKBB data report MAEs in the range of 2.14 to 2.86 years (Tanveer et al., 2023), thus our models lay in the competitive range. The difference between our results and the lowest reported MAE (2.14; Peng et al., 2021) is largely due to the use of performance-enhancing measures such as ensembling, data augmentation, and label binning, which we omitted in order not to jeopardize the generalizability of our model architecture comparison. As the number of available sMRI images in large databases like the UKBB continuous to grow, the SwinT, given its scaling performance in Figure A1, is likely to replace the ResNet as the de facto default deep learning model architecture for brain age prediction.

### 6.2. Potentially different concepts of brain age between model architectures

It would be problematic for brain age research if different model architectures produce different biomarkers, because previous brain age studies’ results regarding clinical utility of BAGs would be confounded by the model architecture. Thus, a critical question remains: do distinct model architectures focus on different concepts of brain age, potentially leading to different biomarkers? Reassuringly, we found no evidence that model architectures consider different concepts of brain age (Section 5.2), mitigating concerns about model architecture confounding in deep brain age studies. Given the conceptually distinct model architectures analyzed, which we consider the most plausible cause of potential variations in brain age concepts, we believe that our results should generalize to related model architectures, such as various CNNs used in previous brain age studies (Peng et al., 2021; Huang et al., 2017; Kolbeinsson et al., 2020).

Beyond that, our results suggest that the clinical utility of BAGs is independent of the deep learning model architecture. Hence, our study provides no reason to consider clinical utility when selecting a model architecture for predicting brain age.

### 6.3. Potentially different concepts of brain age across train runs

Another cause of concern is that random influences such as weight initialization and batch order in training could affect concepts of brain, since many brain age studies’ results base on a single model instance (Bashyam et al., 2020; Cole et al., 2017; Jónsson et al., 2019). Our results alleviate such concern, as the brain age concepts of sViT, ResNet, and SwinT appeared stable across different training runs. Compared to model architecture potentially confounding brain age studies, issues related to random influences are less problematic, because ensembling could be used to account for any variance within a model architecture.

### 6.4. Relation between deep learning model architecture’s accuracy and clinical utility

Our results suggest that clinical utility and deep learning model architecture are unrelated, however, we also found that the noticeably less accurate sViT generated BAGs with very similar clinical utility to BAGs of the more accurate SwinT and sViT, which contrasts with the common belief that more accurate models lead to more useful biomarkers (Hahn et al., 2021; Peng et al., 2021; Cole, 2020; Tanveer et al., 2023; Niu et al., 2020). Along the same lines, previous work has questioned the relation between accuracy and clinical utility: Bashyam et al. (2020) have reported that stopping CNNs’ training before convergence increases biomarker utility; Jirsaraie et al. (2023) have reviewed multimodal brain age studies, including deep and traditional ML models, and have not found a relation between accuracy and clinical utility; Schulz et al. (2024b) have shown that simple linear models, yielded more useful biomarkers than their more accurate deep counterparts. Together, the mentioned evidence in conjunction with our work indicates that optimizing how accurately model architectures predict brain age, is not the right way to fully exploit their potential for generating useful biomarkers. Instead, research in altering the training protocol such as Bashyam et al. (2020) (early stopping) and Schulz et al. (2024b) (overregularization) appears to be more promising.

### 6.5. Questionable construct validity of brain age

In the present study we found no evidence of interaction between model architecture, weight initialization, and batch order and a models’ concepts of brain age. However, Schulz et al. (2024b) reported that reducing a model’s expressivity via overregularisation can indeed yield different concepts of brain age. Such models, while sacrificing age prediction accuracy, seem to provide superior clinical utility. Other work questions whether individual differences in brain age relate to aging effects at all: Vidal-Pineiro et al. (2021) argue that birth-weight and genetic factors have greater impact on BAGs than actual longitudinal brain change. These findings challenge the construct validity of the brain-age gap itself, leading to an increased urgency to develop clearer terminology and methodology to investigate the underlying concepts of brain age learned by machine learning algorithms.

### 6.6. Limitations

We would like to highlight the three important limitations of our study: First, we did not optimize the transformers for prediction accuracy, due to computational constraints. Training a single instance of either transformer model architecture required multiple days on the available GPUs, and the hyperparameter configuration space is vast, making thorough optimization impractical. As a result, the reported accuracies should be viewed as promising lower bounds to optimized accuracies rather than precise estimates. We encourage further work to deep dive in optimizing especially the SwinT’s accuracy in predicting brain age.

Second, we argue that there is most likely no difference in concepts of brain age between different model architectures, however, we are aware that proving such absence of a difference is conceptually hard. That is, it is nearly impossible to cover all conditions (architectures, hyperparameters, demographic factors) for which predictions and prediction errors could indicate differences in concepts of brain age. Nevertheless, we believe that the presented absence of a difference in brain age concepts between model architectures is instructive, because we carefully chose deep learning model architectures based on differences in fundamental design principles, which we consider the most plausible cause of differences in brain age concepts. Also, we did cover a broad set of demographic factors with various neural correlates (Raz et al., 2005; Geng et al., 2006; Gallinat et al., 2006; Dekker et al., 2021; Gómez-Apo et al., 2021), which in its entirety is most likely sensitive to meaningful differences in brain age concepts.

Third, we analyzed predictions and prediction errors as proxies to the underlying concepts of brain age, due to lack of more precise methods, such as reliable explainable artificial intelligence (XAI) methods (Sundararajan et al., 2017; Hooker et al., 2019; Adebayo et al., 2018; Ghorbani et al., 2019; Kindermans et al., 2019; Dombrowski et al., 2019). Such XAI methods (Bach et al., 2015; Selvaraju et al., 2017; Chefer et al., 2021; Ali et al., 2022; Lundberg and Lee, 2017) provide heatmaps indicating which parts of the input have been relevant for ML models’ predictions, which, in theory, seems appropriate to investigate in whether different model architectures learn different brain age concepts. However, in practice, common XAI methods have failed to meet fundamental axioms (Sundararajan et al., 2017), have not been able to beat random relevance assignments (Hooker et al., 2019), have generated heatmaps independent of model parameters and training data (Adebayo et al., 2018), and are susceptible to imperceptible changes in input (Ghorbani et al., 2019; Kindermans et al., 2019; Dombrowski et al., 2019). In light of the mentioned concerns, we believe that it is crucial to validate that XAI methods can reliably explain deep architectures’ predictions in the neuroimaging domain, before their application. To our knowledge, however, explanation methods have not been validated on brain data. This is, because validation of XAI methods often defaults to visual inspection (Doshi-Velez and Kim, 2017), but expectations on explanations in the brain imaging domain are often apriori unknown, or highly difficult to characterize.

### 6.7. Conclusion

In this work, we highlight the possibility of heterogeneity in “concepts of brain age” learned by modern machine learning algorithms.

Reassuringly, we found no indications that deep learning model architectures attend to different concepts of brain age, and hence, it appears unlikely that previous deep brain age studies’ results, for example regarding the clinical utility of BAGs, have been confounded by the model architecture used.

## Appendices

### A. Adapting vision transformers to predict age from 3D sMRI scans

We adapted both SwinT and sViT to operate on 3D input, by dividing input images into 3D image cubes, instead of 2D image patches. Also, we used sinusoidal positional encodings (Vaswani et al., 2017) in the SwinT, in addition to the relative position bias present by default. Sinusoidal positional encodings provide information on an image cubes’ absolute positions in the input image. We anticipated that information on absolute cube position would benefit the model architectures, given that we linearly registered input images to the MNI152 reference space, which leads to image cubes displaying very similar brain regions across subjects. Similarly, sinusoidal positional encodings were employed in the sViT as part of its default setting. Finally, we applied linear regression layers after the transformer-based encoders, to obtain scalar age predictions.

**Table A1:** Hyperparameters for the 3D sViT.

|  |  |
| --- | --- |
| Layers | 6 |
| Heads per layer | 8 |
| Patch size | 16, 16, 16 |
| Embedding size | 1024 |
| MLP size | 2048 |

**Table A2:** Hyperparameters for the 3D SwinT.

|  |  |
| --- | --- |
| SwinT blocks per stage | 2, 2, 6, 2 |
| Attention heads in blocks of each stage | 3, 6, 12, 24 |
| Patch size | 2, 2, 2 |
| Attention window size | 4, 4, 4 |
| Initial embedding size | 48 |
| MLP expansion factor ($\alpha$) | 4 |

### B. Probing how model architectures’ accuracies scale with sample sizes

We investigated how each model architectures’ accuracies relate to the number of training samples, by training 3 instances of sViT, SwinT, and ResNet on different fractions of the training samples (0.2, *n* = 5507; 0.4, *n* =11015; 0.6, *n* =16522; 0.8, *n* =22030). The reduced train sets were generated by iteratively removing the last participants from the original train set. To extrapolate each model architecture’s accuracy beyond our available training sample size, we adopted a power-law relationship between accuracy and the number of training samples, as suggested by Schulz et al. (2024a) (Figure A1). Our analysis indicates that SwinT and sViT will see substantial accuracy improvements with more training samples, whereas the ResNet will experience only marginal gains; the SwinT is expected to surpass ResNet in accuracy at around 25,000 training samples.

**Figure A1:**
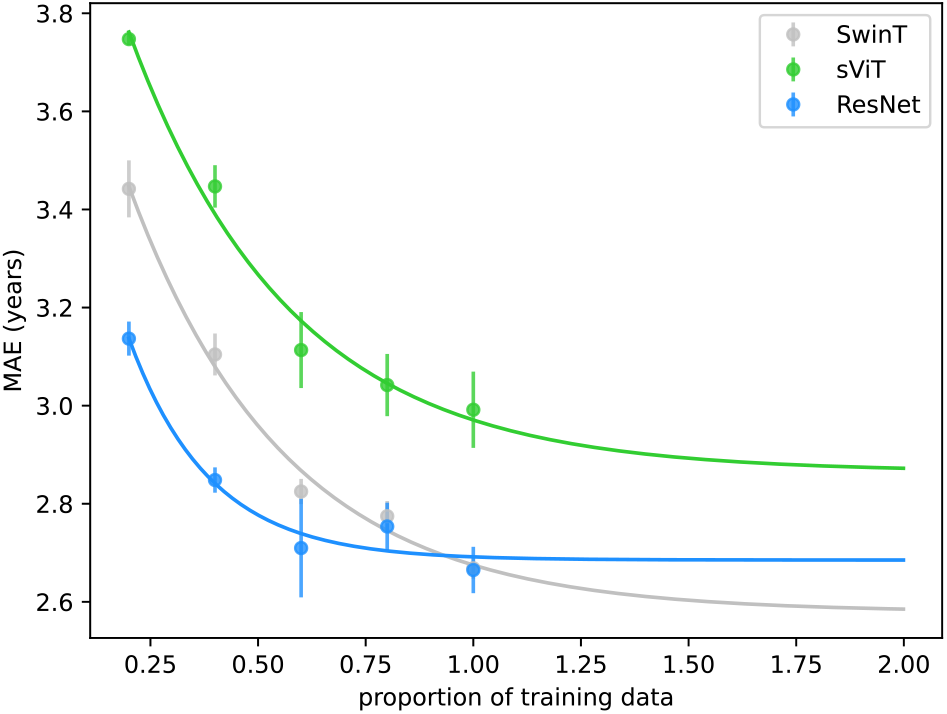
SwinT is highly likely to outperform ResNet with additional training samples We trained multiple instances of each model architecture with gradually decreased training samples and found that accuracies of shifted window transformer (SwinT) and simple vision transformer (sViT) decline stronger compared to the ResNet. Extrapolating each model architecture’s accuracy using power laws (Schulz et al., 2024a) suggests that the SwinT will be more accurate than the ResNet with additional training samples. Uncertainty estimates refer to the SD across model instances.

## Acknowledgments

We thank the UKBB participants for their voluntary commitment and the UKBB team for their work in collecting, processing, and disseminating these data for analysis. Research was conducted using the UKBB resource under project-ID 33073. Computation has been performed on the HPC for Research cluster of the Berlin Institute of Health. The project was funded by the Deutsche Forschungsgemeinschaft (DFG, German Research Foundation) under project-ID 414984028 - CRC 1404 and the Brain & Behavior Research Foundation (NARSAD young investigator grant). K.R. was additionally supported by the DFG (389563835, 402170461 - TRR 265, 459422098 - RU 5363, and 442075332 - RU 5187), and a DMSG research award.

## Author contributions

**Nys Tjade Siegel**: Data curation, Methodology, Software, Formal analysis, Visualization, Writing - Original Draft. **Dagmar Kainmueller**: Formal analysis, Methodology, Writing - Review & Editing. **Fatma Deniz**: Formal analysis, Methodology, Writing - Review & Editing. **Kerstin Ritter**: Conceptualization, Methodology, Formal analysis, Writing - Review & Editing, Funding acquisition. **Marc-Andre Schulz**: Conceptualization, Methodology, Software, Formal analysis, Writing - Original Draft, Writing - Review & Editing, Supervision.

## Declaration of Generative AI and AI-assisted technologies in the writing process

During the preparation of this work the author(s) used Microsoft Copilot, GPT-3.5 and GPT-4 in order to improve readability and language. After using this tool/service, the author(s) reviewed and edited the content as needed and take(s) full responsibility for the content of the publication.

## Supplementary Material

**Table S1:** Mappings from diseases (first occurrence dates) to UK Biobank field numbers.

| **Disease** | **Field number** | **Instance** | **Array index** |
| --- | --- | --- | --- |
| Parkinson's disease | 131022 | 0 | 0 |
| Multiple sclerosis | 131042 | 0 | 0 |
| Epilepsy | 131048 | 0 | 0 |
| Alcohol use disorder | 130854 | 0 | 0 |
| Bipolar affective disorder | 130892 | 0 | 0 |
| Schizophrenia | 130874 | 0 | 0 |
| Schizotypal disorder | 130876 | 0 | 0 |
| Persistent delusional disorders | 130878 | 0 | 0 |
| Acute and transient psychotic disorders | 130880 | 0 | 0 |
| Induced delusional disorder | 130882 | 0 | 0 |
| Schizoaffective disorder | 130884 | 0 | 0 |
| Other nonorganic psychotic disorders | 130886 | 0 | 0 |
| Unspecified nonorganic psychosis | 130888 | 0 | 0 |

**Table S2:**
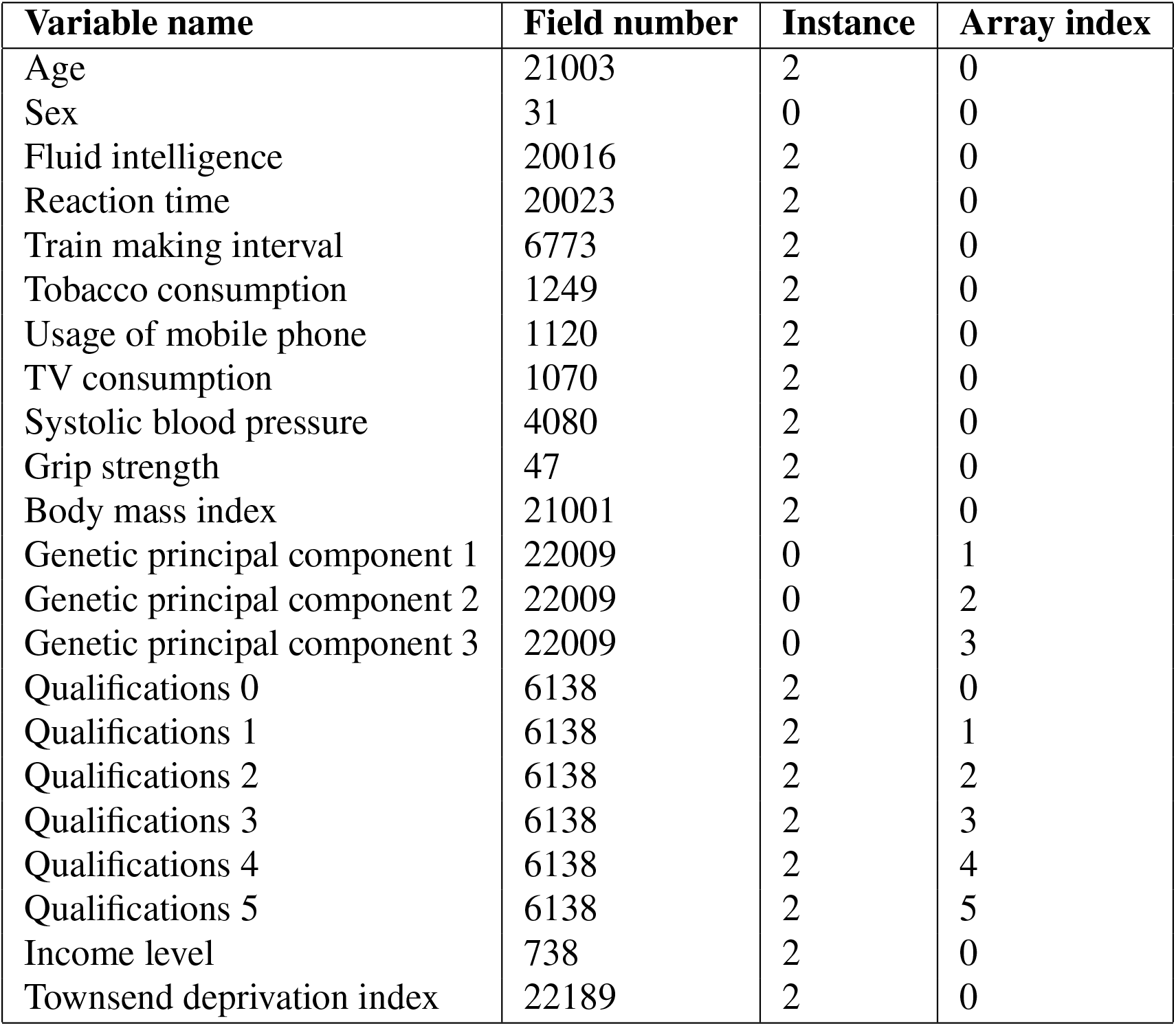
Mappings from variable names to UK Biobank field numbers.

## Footnotes

1 Psychotic disorders refer to schizophrenia, schizotypal and delusional disorders (ICD-10 codes F20 to F29). They are treated as a single group in our analysis due to the impractically small sample sizes (*n* < 33) when treated separately.

2 https://github.com/kenshohara/3D-ResNets-PyTorch

3 https://github.com/lucidrains/vit-pytorch

